# Coping-Style Behavior Identified by a Survey of Parent-of-Origin Effects in the Rat

**DOI:** 10.1101/351155

**Authors:** Carme Mont, Polinka Hernandez Pilego, Toni Cañete, Ignasi Oliveras, Cristóbal Río-Álamos, Gloria Blázquez, Regina López-Aumatell, Esther Martínez-Membrives, Adolf Tobeña, Jonathan Flint, Alberto Fernández-Teruel, Richard Mott

**Affiliations:** Wellcome Trust Centre for Human Genetics, University of Oxford, OX3 7BN, UK; Medical Psychology Unit, Department of Psychiatry & Forensic Medicine, Institute of Neurosciences, Universidad Autónoma de Barcelona, 08193-Bellaterra, Barcelona, Spain; Fundació Bosch i Gimpera, Parc Científic de Barcelona, 08028, Spain; Universitat Pompeu Fabra, Department of Experimental and Health Sciences, 08003 Barcelona, Spain; Center for Neurobehavioral Genetics, University of California Los Angeles, 695 Charles E. Young Drive South, Los Angeles, California USA; Genetics Institute, University College London, Gower St, London WC1E 6BT, UK

## Abstract

We develop theory, based on earlier work, to partition heritability into a component due to a combination of parent of origin, maternal, paternal and shared environment, and another component that estimates classical additive genetic variance. We then investigate the effects on heritability of the parental origin of alleles in outbred heterogeneous stock rats across 199 complex traits. Parent-of-origin-like heritability was on average 2.7-fold larger than classical additive heritability. Among the phenotypes with the most enhanced parent-of-origin heritability were 10 coping style behaviors, with average 3.2-fold heritability enrichment. To confirm these findings on coping behaviour, and to eliminate the possibility that the parent of origin effects are due to confounding with shared environment, we performed a reciprocal F1 cross between the behaviourally divergent RHA and RLA rat strains. We observed parent-of-origin effects on F1 rat anxiety/coping-related behavior in the Elevated Zero Maze test. Our results are the first to assess genetic parent-of-origin effects in rats, and confirm earlier findings in mice that such effects influence mammalian coping and impulsive behavior.

## Introduction

Parental genotypes do not necessarily make equal contributions to the phenotypes of their offspring ^1^ but the magnitude of these parent of origin-effects (PoEs) is largely unexplored ^2^. In placental mammals, around 100 transcriptional units are imprinted^3^, i. e. mono-allelically expressed in a parent-of-origin specific manner. The identities of these genes and the parental origin of the silenced allele depend on the species, tissue and developmental stage, with a core set of imprinted genes common to most species. In humans, disruption of specific imprinted genes cause developmental syndromes ^4^, and while the overall impact of PoE on human complex traits is less clear a handful of examples are known; for example PoE at the imprinted *DLK1-MEG3* locus associates with type I diabetes ^5^.

More is understood about PoE in mice. The phenotypic impact of PoE, as assessed in controlled reciprocal crosses and in populations of mice in which the parental origin of alleles can be determined, suggests PoE can be considerable. In ^6^, quantitative trait loci (QTLs) for growth were mapped in an advanced intercross of mice in which offspring and their parents were genotyped: a significant fraction of the phenotypic variance attributable to QTLs was due to PoE. In ^7^, heritability for 100 diverse phenotypes in a heterogeneous stock of mice, again with genotyped parents, was estimated based on genomewide genetic similarity, and partitioned into variance components representing PoE and non-PoE. The PoE component was on average about twice the non-PoE component. Similarly, phenotyping of the offspring of reciprocal crosses between inbred strains of mice have revealed PoE on growth^7^, behavior^8^ and gene expression, in addition to detailed maps of mono-allelic expression as a function of developmental stage and tissue ^9^.

PoE is widespread in many other species, although reports are less comprehensive. Cross-bred cattle harbour PoE QTLs^10^. However, only weak evidence for PoE was observed in poultry^11^. PoE (but not necessarily imprinting) is present in other organisms such as *Drosophila melanogaster* ^12^ and *Caenorhabditis elegans* ^13^, despite the absence of DNA methylation in either species. PoE and imprinting affect embryogenesis in flowering plants ^14^, and extensive epigenetic effects on flowering and other phenotypes occur in *Arabidopsis thaliana* ^15^.

However, there are potential issues of confounding when estimating PoE in populations, and which might lead to other effects such as shared environment or dominance being mis-interpreted as PoE. Both of the family-based mouse studies of PoE ^6,7^ included siblings that are also littermates. Siblings share more alleles by descent than non-siblings, and this excess is driven by alleles shared by common parents ^7^. Thus shared environment (eg maternal effects and cage effects) potentially inflates the variance attributed to PoE. However, it should be noted that cage effects were first removed in the calculation of heritability in ^7^. Dominance effects can also be confounded with PoE ^11^. Carefully designed and replicated reciprocal cross experiments can be used to eliminate these possibilities, and have confirmed that in general PoE are real, although undoubtedly dominance and environment also play significant roles in the architecture of complex traits.

Furthermore, while the studies ^6,7^ associate genetic differences with PoE, they do not suggest a mechanism, except that the PoE QTLs they identified did not typically overlap with known imprinted genes (most of which cluster into about 13 imprinted loci^16^). Thus the plausible assumption that only variation *in cis* at imprinted loci causes phenotypic PoE - notwithstanding syndromes in humans due to mutations near imprinted genes^4^ - is not true. Indeed, transcriptome analysis of reciprocal crosses in ^7^ suggests that natural genetic variation *in trans* perturbs mono-allelic expression of imprinted genes to cause phenotypic PoE. Similarly, physiological PoE ^7,17,18^ have been observed in reciprocal crosses between knockouts (KO) of non-imprinted genes and wild-type mice with isogenic backgrounds. Both phenotypic PoE and parental effects were observed, often sex-specific.

It is therefore important to ask whether broadly similar phenomena occur in species other than mice. In rats, prenatal glucocorticoid overexposure generates multigenerational epigenetic effects on weight and on DNA methylation, transmitted through both paternal and maternal pathways ^19,20^. Similarly, a high-fat diet generates metabolic PoE ^21^. These epigenetic effects suggest it would be worthwhile to assess the impact of genetic variation on PoE in rats. Finally, rats exhibit more complex behaviours than mice, particularly coping-style behaviors, so it is especially important to consider PoE in the context of rat behavior.

Here, we use two experimental rat resources. We start with data from a rat heterogeneous stock previously analysed to map classical additive QTLs ^22^. We ask whether heritability of complex traits shows a similar breakdown to that seen in mice. We derive a new parameterization of the variance components model used to partition heritability into PoE-like effects (true PoE, maternal, paternal and shared environment) and non-PoE, which has some advantages of interpretation to that presented previously in ^7^.

We then focus on rat behavior in greater detail, to determine whether PoE influence coping style behaviors in a reciprocal cross between RLA-I and RHA-I Roman rat strains ^13^. These, in addition to the HS rat stock, are ideal for studying the genetic basis of two-way-shuttle box-avoidance acquisition, classically conditioned fear/freezing and unconditioned anxiety in the elevated plus-maze and zero-maze ^22–24^.

As a population, HS rats have an emotional profile similar to that of RLA-I rats, which is characterised by enhanced anxious and fearful responses in conditioned and unconditioned anxiety tests such as the elevated zero maze, classically-conditioned freezing and the acquisition of two-way active avoidance in the shuttle box ^25,26^; such responses are classified as reactive (“passive”) coping styles. In contrast, a pro-active coping style comprises active responses directed to address an aversive situation, either to confront stressful stimuli or to increase the distance between the stressful stimuli and the individual ^27^. Pro-active coping styles, which are characteristic of RHA-I rats, include increased exploratory behavior ^28–31^, aggression ^32,33^ and struggling ^29,34,35^. The reactive coping style involves inactivity and disengagement when in an aversive situation, only reacting if absolutely necessary. The reactive coping style has been associated with non-aggressive behavior ^35,36^, absence of responses ^29,33,37^, and immobility and extended latency when facing novel/stressful situations ^29,30,34,38^. Impulsivity is closely related to coping ^39^, and genetic perturbation of specific imprinted genes leads to changes in impulsive behavior in mouse reciprocal crosses^40–42^. We therefore investigated the extent to which rat behaviour is influenced by PoE in general, and how rats and mice compare in their PoE responses to coping and impulsivity.

## Materials and Methods

### Parent-of-origin heritability in the HS rats

The HS rats used in this experiment are a subset of those reported in ^22,43^. Genotypic data was available for 798 individuals, for which both parents were genotyped (198 parents) at 265,551 SNPs using the RATDIV SNP array. Phenotypes for 199 traits were available for subsets ranging from to 205 to 617 individuals (Supplemental Table S1). Estimates of heritability were obtained according to ^7^ (Methods). We used the R ^44^ package happy.hbrem ^45^ to compute phased haplotype probabilities, from which kinship matrices were computed and analyzed. Heritability was computed using GCTA ^46^ applied to these Genetic Relationship Matrices (GRM).

The variance decomposition used in ^7^ separates the total phenotypic variance *σ*^2^ into three parts, namely 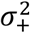 associated with allele-sharing from parents of the same sex (and interpretable as due in part to parent-of-origin, maternal and paternal effects, and potentially shared environment when siblings are co-housed), 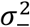 associated with allele-sharing from parents of the opposite sex (i.e. the additive genetic variance not attributable to PoE), and 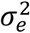, the environmental variance. In the Methods we show this decomposition can be written in a mathematically equivalent way that makes its biological interpretation clearer. Another decomposition of heritability into maternal and paternal components is described in ^11^.

First note that each genotype of a diploid individual can be split into maternally and paternally inherited components. In a population sample of *n* individuals genotyped at *l* loci, these can be represented as a pair of *n* × *l* matrices *M,P* respectively, such that the *i, k* element is the appropriately normalised maternally or paternally inherited allele of individual *i* and locus *k*. The total genotype dosage matrix *G* = *M* + *P*. If the genotypes are all biallelic SNPs, and the SNP *k* is in Hardy-Weinberg equilibrium with allele frequency *π*_*k*_ then the un-normalised genotype dosages decompose into their maternal and paternal components as *g*_*ik*_ = *m*_*ik*_ + *p*_*ik*_. The normalized genotype dosage matrix *G_ik_* decomposes into normalized maternal and paternal components thus:

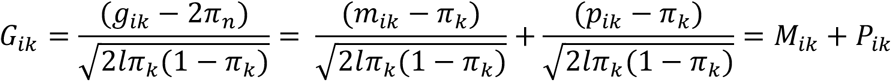

If the genetic markers are multi-allelic, or represent local haplotypes, then the definitions are slightly different. In ^7^ and in this study we use matrices defined in terms of the probability that an individual carries particular maternally inherited and paternally inherited haplotypes at a given locus. The formulae are given in ^7^ and are not repeated here (the decomposition for SNPs given here is new). Earlier work on partitioning kinship matrices into maternal and paternal components is described in ^11,47^. The important point is that matrices *M*,*P* can be constructed, representing maternal and paternal genotype or haplotype contributions as required.

The additive GRM *K,* of dimension *n* × *n*, that is routinely used to estimate heritability is usually defined as^46^:

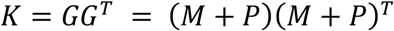

(where the superscript *T* indicates matrix transposition). It can be partitioned thus:

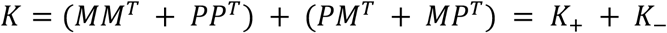

Note that by construction the expected values of the diagonal elements of the standard symmetric GRM *K* are all unity, and the off-diagonal elements *K*_*ij*_ are the correlations between individuals *i,j* based on their normalized genotypes. The matrix *K*_+*ij*_ is the component of the genetic relationship between *i*,*j* that is attributable to allele-sharing from parents of the same sex. Thus if *i*,*j* are siblings we mean inheritance from the same individual parent (i.e. identity by descent), otherwise inheritance of the identical allele but from distinct individuals of the same sex (i.e. identity by state and inherited from a parent of the same sex). The matrix *K*_−*ij*_ is the component of allele-sharing in which the parents transmitting the shared allele are of opposite sex. Since the corresponding diagonal elements of *K*_+_, *K*_−_ are all positive and sum to unity in expectation, individually they are less than unity.

The variance matrix of a Multivariate-Normally distributed phenotype *z* is modeled as a sum of these matrices, plus a diagonal matrix of uncorrelated environmental effects:

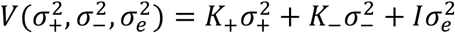

Under the null hypothesis that there are no PoE then 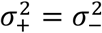 and the variance matrix collapses to the simpler and more familiar form

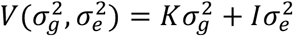

where 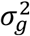 is the usual additive genetic variance.

### Reparameterisation

It is desirable that each component of the variance decomposition should itself be a variance component, that is both V and each of the individual matrices in the decomposition should be positive semidefinite (PSD). By construction, both *K* and *K*_+_, are positive-semidefinite (PSD) matrices since for any vector *z*,

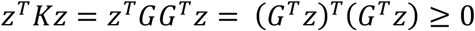

and

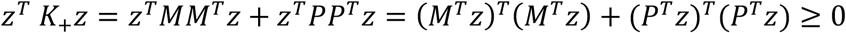

However, *K*_−_ is not of the same form and is not necessarily PSD. We provide here a reparameterisation that solves this difficulty and provides an alternative interpretation of the decomposition. First note that the log-likelihood of the data satisfies

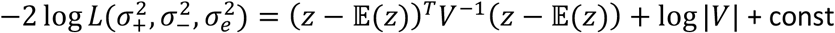

which depends on 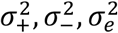 solely through *V,* so any re-parameterisation that preserves *V* will also preserve the likelihood, and so have equivalent maximum-likelihood estimates. The diagonal entries are less than unity and 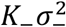 is not PSD. However, if we define 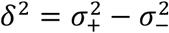 we can reparameterise the variance as

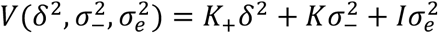

so that each component is PSD. Under the null hypothesis of no PoE, *δ*^2^ = 0 and the variance becomes that of the standard additive genetic model. The valid parameter space for this variance component is *δ*^2^ ≥ 0, i.e. 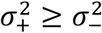. In the region *δ*^2^ > 0 the likelihoods are equivalent under both parameterisations so the maximum likelihood estimates (MLEs) satisfy 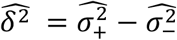. However, if 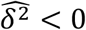 then 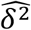 will be set to zero (i.e. moved to the boundary) and the two parameterisations will report different MLEs. These observations also apply to the heritabilities 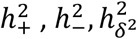 associated with 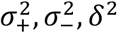 respectively, each computed by dividing by the total phenotypic variance.

This reparameterization also shows, irrespective of the truth of the null hypothesis, that 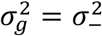. In other words, 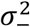 is an estimator of the additive heritability 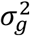 that would be estimated in population samples without any first-degree relatives (e.g. in genetic association studies). Thus, even in a population sample containing siblings, provided their parents are not related, the variance matrix *K*_−_ = *MP*^*T*^ + *PM*^*T*^ is an unbiased estimator of the matrix 0.5*K* that would be obtained from a sample of unrelated individuals, in the sense that

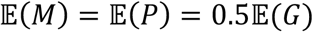

so that, because *M*, *P* are independent,

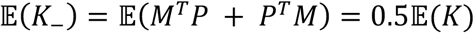

Thus 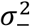 estimated from related individuals equals the additive heritability that would have been observed in a sample of unrelated individuals. In contrast, 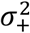 is the additive heritability, possibly inflated by family relatedness should the null hypothesis be false. The excess heritability *δ*^2^ can be caused by PoE, but may also be confounded by shared environment, for example maternal effects.

One can then make inferences about genetic architecture via hypothesis tests on the m.l.e.s of 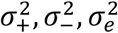 using the information matrix to generate variance and covariance estimates. It is worth noting that the likelihood is well defined whenever the variance matrix *V* is positive definite (PD), which is the case even for small but negative values of *δ*^2^. Since 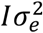 is always positive definite, a sufficient condition that *V* is positive definite is that the genetic component 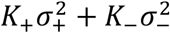 also be positive definite. Using the fact that both *K* = *K*_+_ + *K*_−_ and *K*_+_ are positive definite, and that

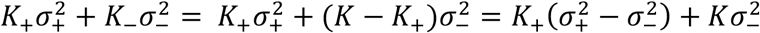

it follows that this is positive definite whenever 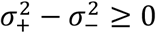. Thus the matrix *V* is guaranteed to be positive definite whenever 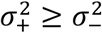, which is precisely the region of interest in a PoE analysis. In fact, provided 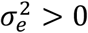, the region for which *V* is positive definite will also include some 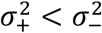, the boundary depending on *K*_+_,*K*_−_ and on the phenotype vector in question. However, there do not appear to be biologically realistic scenarios for which 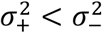. Under the null hypothesis that there is no PoE then 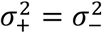. This can be thought of as being an edge of the parameter space.

Finally, it is also worth remarking that partitioning the variance into 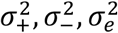 differs from spatial partitioning, such as when each chromosome has a unique GRM and heritability ^46^, or when the genome is divided according to feature annotation^48^. It more closely resembles the approach in ^49^, in which a GRM approximating the IBD degree of genetic relatedness expected due to the pedigree is effectively subtracted from *K* to generate approximate analogues of the matrices *K*,*K*_+_, used here. The details of the matrix construction are different in ^49^, however.

### RHA/RLA Roman Rats F1 reciprocal cross

The Wistar-derived outbred sublines of Roman High-(RHA/Verh) and Low-(RLA/Verh) Avoidance rats have been genetically selected since 1972 because of their good (RHA/Verh) versus extremely poor (RLA/Verh) acquisition of two-way, active avoidance^29,50^. The inbred strains RHA-I and RLA-I, derived from those two lines, have been maintained at our laboratory in the Autonomous University of Barcelona since 1996^50,51^.

Reciprocal crosses between the 60^th^ generation of RHA-I and the RLA-I inbred rat strains (hereafter RHA and RLA, respectively) were set up with the following configuration: female RHA/male RLA (fRHA/mRLA) and female RLA/male RHA (fRLA/mRHA). The breeding pairs were housed together and separated once pregnancy was confirmed. Animals were selected from the F1 crosses, comprising 34 fRHA/mRLA pups (19 females and 15 males) and 37 fRLA/mRHA pups (22 females and 15 males). Pups were weaned and caged in pairs of siblings in macrolon cages (50 cm × 25 cm × 14 cm). They were maintained with food and tap water available ad libitum, under conditions of controlled temperature (22°C ± 2°C; 50% - 70% humidity) and a 12-h light/12-h dark cycle (lights on at 08:00 h). Behavioral testing commenced at 8-10 weeks of age, with one-week separation between the two tests. Experiments were performed at the Medical Psychology Unit, Department of Psychiatry & Forensic Medicine, Autonomous University of Barcelona, Spain, between 09:00 and 19:00h in accordance with the Spanish legislation on “Protection of Animals Used for Experimental and Other Scientific Purposes” and the European Communities Council Directive (86/609/EEC) on this subject.

### Elevated Zero Maze

The maze comprised an annular platform (105 cm diameter; 10 cm in width) made of black plywood and elevated to 65 cm above the ground level. It had two open sections (quadrants) and two enclosed ones (with walls 40 cm in height). The subject was placed in an enclosed section facing the wall. The apparatus was situated in a black testing room, dimly illuminated with red fluorescent light, and the behavior was videotaped and measured outside the testing room by an expert observer who was blind to the cross condition. Latency to enter into an open section, time spent in the open sections, number of entries in the open sections, number of head dips (through the edge of the open sections), number of line crossings and number of stretched attend postures (SAP, from a closed to an open section of the maze) were measured for 5 min. All these measures (except “line crossings”, which are an index of overall locomotor activity), have been pharmacologically and behaviorally validated as anxiety-related parameters ^25,52,53^.

### Two-way active shuttle-box avoidance acquisition

Active avoidance acquisition sessions were performed in three identical shuttle boxes (Letica Instruments), each one placed in independent sound-attenuating boxes consisting of two equal-sized compartments (25 × 25 × 28 cm) connected by an opening (8 × 10 cm) and illuminated by a dim fluorescent light (<50 lux). Rats were allowed a 4-min period of familiarization to the box. Immediately after that period, a 40-trial session/rat was administered, each trial consisting of a 10-sec CS (conditioned stimulus; 2400 Hz, 63-dB tone plus a 7-W small light) followed after termination by a 20-sec US (unconditioned stimulus; scrambled 0.7-mA foot shock) delivered through ^53^the grid floor. Crossings to the other compartment during the CS (avoidances) or US (escapes) switched off the stimuli and were followed by a 60-sec inter-trial interval. Avoidances, escapes, latency to escape/avoid, inter-trial crossings (crossing to the other compartment during the inter-trial period) and context-freezing (complete and rigid immobility-except for breathing movements-during the first 5 inter-trial intervals of the avoidance training session) measurements were recorded. Freezing was measured by an expert observer who was blind to the cross direction.

These parameters (in particular, avoidances, latency to escape/avoid, intertrial crossings and context-conditioned freezing) have been pharmacologically and behaviorally validated as anxiety/fear-and coping style-related measures ^26,29,43,50,54,55^ The data was detected and loaded into a computer automatically, except the context-freezing which was measured by a researcher during the first five inter-trial intervals.

### Statistical Analysis of behavior

F1 rat phenotypes were analysed using R. First, a principal components analysis identified a family with extreme values and three subjects of the family were eliminated from the sample for all the measures. The remaining behavioral data were analysed by the R function lmer() in the R package lme4 ^56^. Analyses fitted family (defined as the concatenation of maternal and paternal id) as a random effect and parental origin (encoded by the identity of the maternal strain, mstrain), plus sex as fixed effects (Supplemental Tables S2,3). For example the trait freezing5 would be modeled in R as

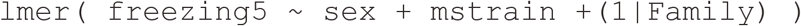

Significance of fixed effects was determined by analysis of deviance with the R command anova(), using the chi-squared distribution on 1 degree of to compute P-values of the PoE effect (ie whether the fixed effect parameter estimate associated with mstrain was significantly different from zero).

## Results

### Heritability and parent of origin effects in HS rats

Following^7^, we define a PoE as a difference in the phenotypic effect due to an HS progenitor haplotype, depending on whether it originated in the previous generation from the father or mother. To do this, we compute the phased probabilities that an animal inherits the ancestral founder haplotype from each parent, and then use these probabilities to partition the kinship by PoE.

Parental genotypes were available for 798 HS rats, allowing genotypes to be phased with respect to parental origin. A list of the phenotypes and number of rats phenotyped for each trait is described in Supplemental Table S1. Using 265,551 single nucleotide polymorphisms (SNPs) to estimate founder haplotypes at each locus and individual, we derived a genetic relationship matrix which was partitioned into components representing allele sharing from parents of the same sex *K*^+^, and from parents of the opposite sex, *K*^−^. Figure 1 plots the distributions of the entries of these matrices, and shows that, consistent with previous work in HS mice, in *K*^+^ (Figure 1A) the siblings have distinctly more allele-sharing than non-siblings. In *K*^−^ (Figure 1B) the distributions are closer together, but interestingly - and in contrast to HS mice - they are still separable. Figure 1B shows that rats selected for mating were more closely related than would be expected if they were chosen at random. If parents are unrelated then the expected distribution of *K*^−^ for siblings and non-siblings should be equal, as was observed for HS mice^7^.

**Figure 1.**
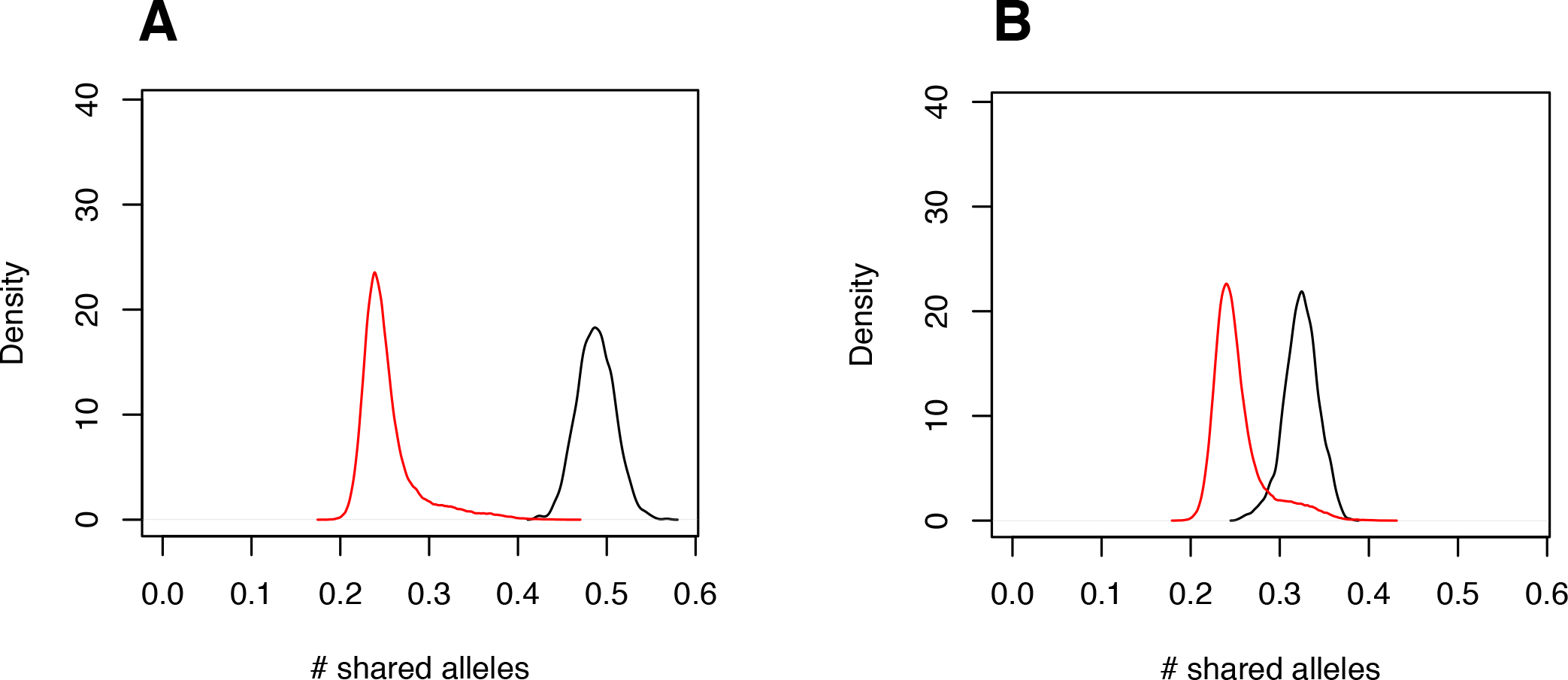
Frequency distributions of the off-diagonal elements of the Genetic Relationship Matrices *K*_+_ (A) and *K*_−_ (B) in HS rats. In each plot, the distributions of siblings (black) and non-siblings are shown separately. x-axis shows the number of shared alleles, y access is the smoothed density.

**Table 1.**
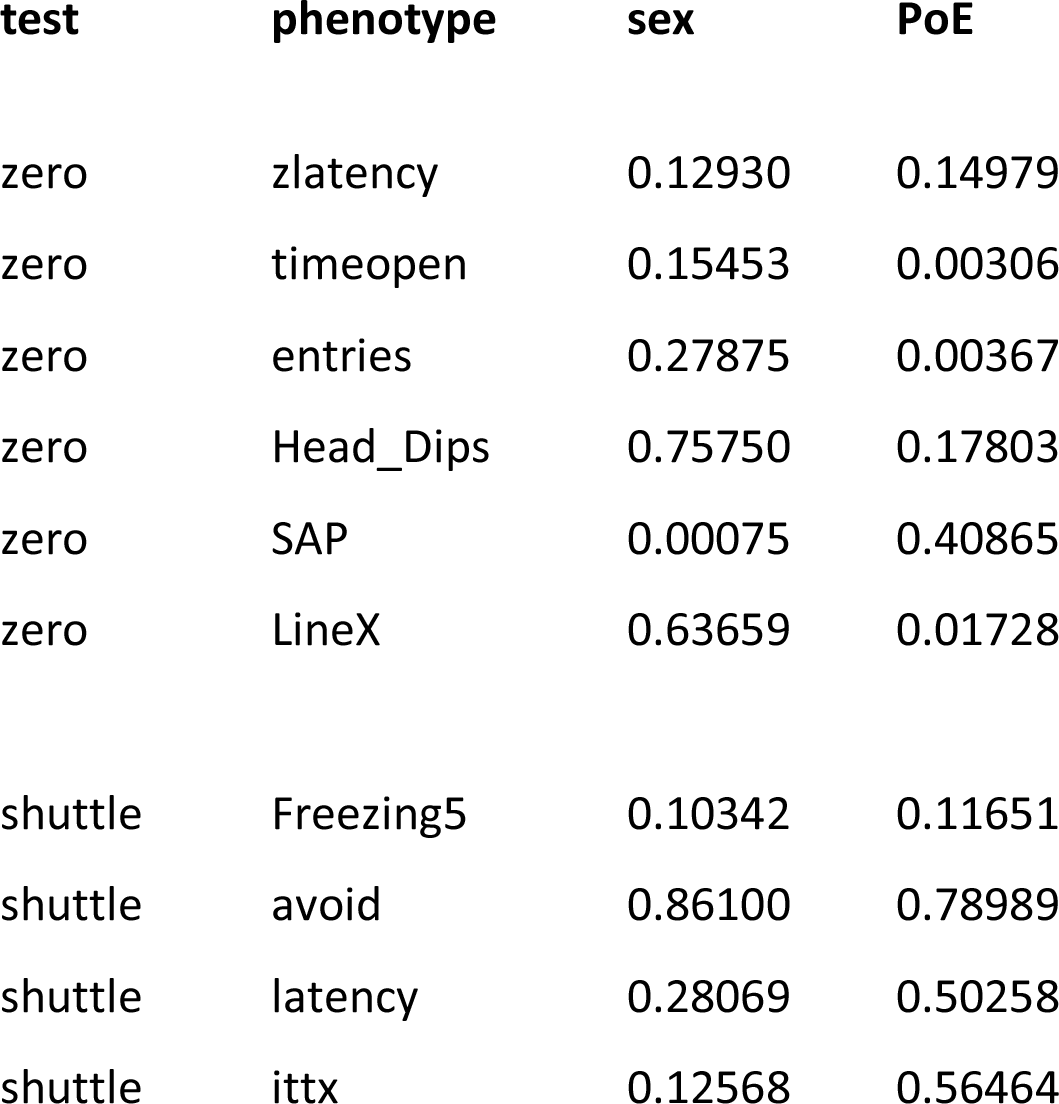
P-values for the effects of sex and parent-of-origin (PoE) for coping-style behavior as measured in 68 rats from a reciprocal cross of RHA and FHA, in the elevated zero-maze (zero) and shuttle box (shuttle). Phenotypes are: zlatency (time/s to first entry into the open sections of the zero maze), timeopen (time/s spent in open sections of the maze), entries (number of entries into maze open sections), Head_Dips (number of head dips through the edge of the open sections of the maze), SAP (number of stretch attend postures from a closed to an open section of the maze), LineX (number of line crossings in the elevated zero maze), Freezing5 (context freezing, i.e. time spent freezing during the first 5 inter-trial intervals of the two-way avoidance-shuttle box-training session), avoid (number of avoidances during the 40-trial avoidance session), latency (time/s to escape, averaged for the 40 training trials of the avoidance session), ittx (inter trial crossings during the forty 60-min inter-trial intervals of the avoidance session). P-values are calculated from a mixed-model analysis of variance.

| test | phenotype | sex | PoE |
| --- | --- | --- | --- |
| zero | zlatency | 0.12930 | 0.14979 |
| zero | timeopen | 0.15453 | 0.00306 |
| zero | entries | 0.27875 | 0.00367 |
| zero | Head_Dips | 0.75750 | 0.17803 |
| zero | SAP | 0.00075 | 0.40865 |
| zero | LineX | 0.63659 | 0.01728 |
| shuttle | Freezing5 | 0.10342 | 0.11651 |
| shuttle | avoid | 0.86100 | 0.78989 |
| shuttle | latency | 0.28069 | 0.50258 |
| shuttle | ittx | 0.12568 | 0.56464 |

Next, for each of the 199 phenotypes we used a mixed model ^46^ to estimate the fraction of phenotypic variation attributable to each component of inheritance 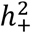 and 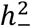. If the parent of origin makes no contribution then we expect 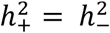. In 86% of the traits examined (172 out of 199), 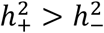 (Figure. 2A). The medians are 0.474 and 0.155 respectively with a median ratio 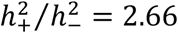. The average standard errors are 0.133 for 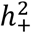 and 0.172 for 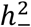. The heritability attributable to 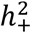 and 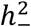 and the total heritability for each of the phenotypes, is detailed in Supplementary Table S1. The numbers of rats for many phenotypes was too small to attempt to map PoE QTLs.

**Figure 2.**
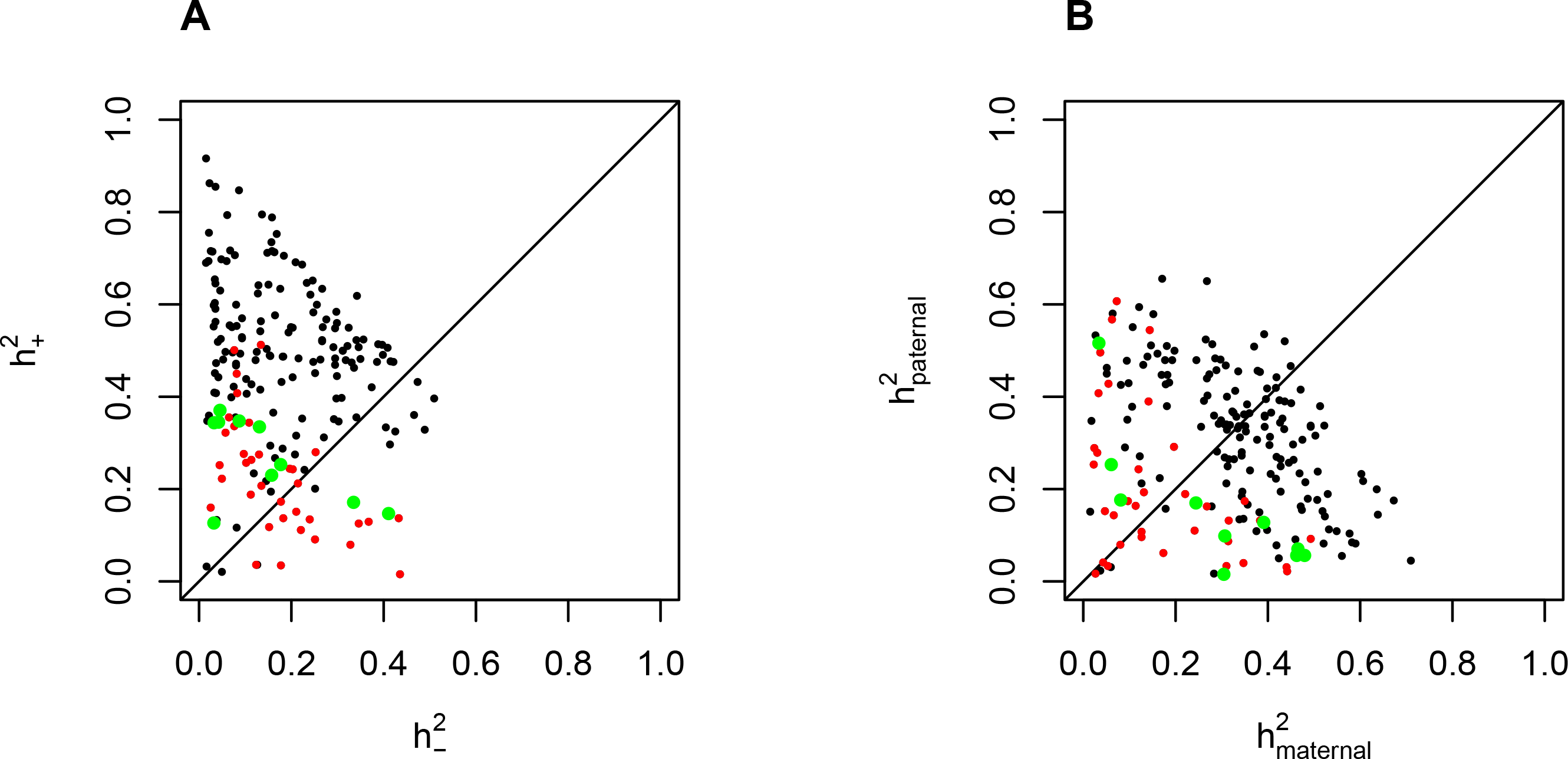
(A) Heritability of 199 traits partitioned into components 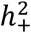 (y-axis) and 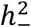 (x-axis) associated respectively with PoE and non-PoE. Each dot represents one trait. Coping-Style traits are in green, other behavioral traits are in red. (B) Heritability partitioned into maternal and paternal components, color-coded as in (A).

In the Methods we show this model for PoE and non-PoE heritability can be re-parameterised in terms of 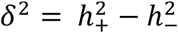 and 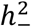. The reparameterisation makes biological interpretation more straightforward: *δ*^2^ is the additional genetic variance explained by PoE and 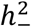 is what would be observed in a sample of unrelated individuals whose parents were also unrelated; in other words 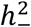 is an estimate of the classical additive heritability.

Covariates such as maternal effects can be confounded with PoE ^57^. Therefore we examined whether the phenotypes showed a higher maternal heritability. For all the 199 traits, the median maternal heritability is 0.323 with a standard error of 0.165, and 0.295 for the paternal heritability with a standard error of 0.164. The median ratio of maternal heritability/paternal heritability = 1.08 (Figure 2B, Supplemental Table S1). We thus find little evidence that maternal effects play a larger role than paternal effects. A recent study of paternally transmitted epigenetic effects on rats exposed to glucocorticoids supports this general observation ^20^.

### Parent of origin effects contribute to coping-style behavioral phenotypes

We analysed 45 behavioral measures, including phenotypes from the elevated zero maze, the novel-cage activity test, context-conditioned freezing and the two-way active avoidance test. We found that these measures had lower than average 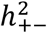 values (Figure 2A). In 57% of the behavioral traits examined (26 out of 45 measures), 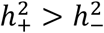. The average standard errors of the heritability estimates were 0.122 and 0. 171 respectively. The medians were 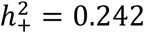 and 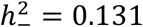 with a median ratio of 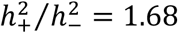.

We next examined the parent of origin effects in ten coping-style behaviors (listed in Supplementary Table S1), In eight out of the ten measures 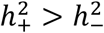 (Figure 2A). The medians are 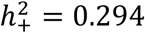 and 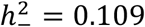 with a median ratio of 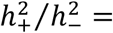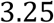 and standard errors of 0.137 and 0.193 respectively. Thus in general behavioural traits show less evidence of PoE, except for coping-style behaviors which exhibit stronger PoE than do most other traits.

### Reciprocal crosses between RHA-RLA strains shows differences in coping style depending on parental origin

We verified the findings on coping style behaviors in the HS by testing for PoE in a reciprocal cross. As noted above, RHA and RLA are inbred rat strains derived from the Wistar stock of rats in an experiment that selected for differences in coping behavior (i.e. acquisition of the two-way active avoidance response). RLA rats show increased stress-induced endocrine responses, enhanced anxiety/fearfulness and passive/reactive coping style in a variety of unconditioned behavioral variables and tests, compared to RHA ^29,50,54^. We therefore chose these strains for behavioral testing in preference to founders of the HS in order to increase the power to detect parent of origin effects on coping behavior.

We made reciprocal F1 crosses (female RHA x male RLA (fRHA/mRLA) or female RLA x male RHA (fRLA/mRHA) and produced 137 offspring in total. The mRHA/fRLA was more fertile than the other cross (103 pups vs. 34) so we phenotyped offspring at random to balance the numbers of families, offspring sex and totals for the two cross directions. We measured behavioral responses in the elevated zero maze and the two-way active avoidance in the shuttle box. In total phenotypes from 68 offspring from 19 families were analysed.

All rats within a family have the same parental origin of alleles. Therefore, to eliminate possible confounding between families and PoE, we fitted a mixed model in which membership across the 19 families was treated as a random effect, while cross direction (parent of origin) and sex were fixed effects. Controlling for family membership in this way enables the testing for parent of origin effects and reduces the possibility of a false positive result due to random differences between familial environments.

In the elevated zero maze, fRHA/mRLA rats enter the open section earlier, make a higher number of entries, explore more quadrants of the maze, and remain in the open section for longer, showing an active coping style similar as the RHA rat strain (Figure 3A-C). fRLA/mRHA rats show the opposite behavioral response, characteristic of a reactive coping style and similar to the RLA strain. For these measures, the behavioral profile of the fRHA/mRLA and fRLA/mRHA rats is more similar to their maternal strain than to their paternal strain.

**Figure 3.**
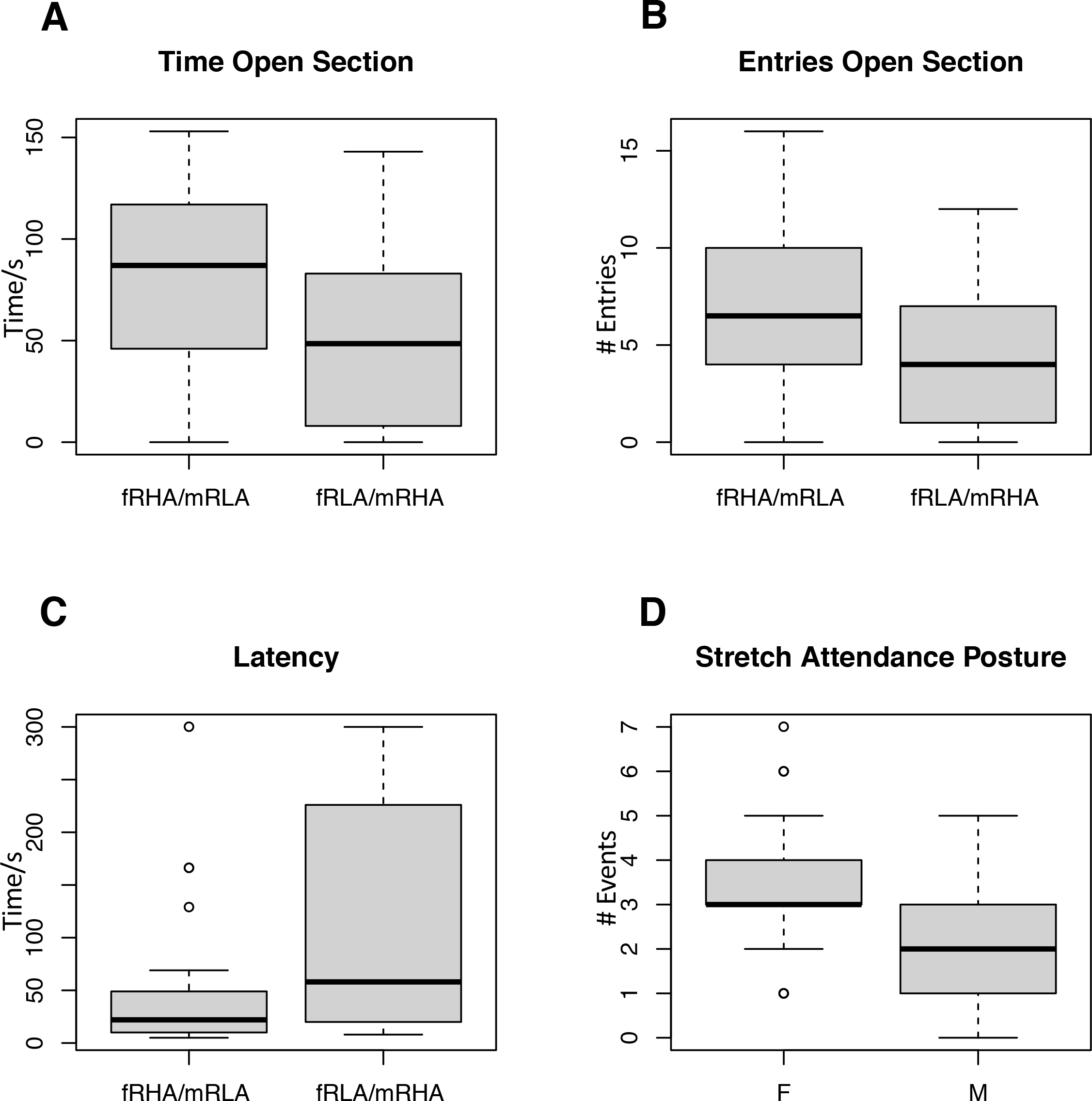
Box-and-whisker plots showing parent-of-origin effects on three behaviors (A-C) and gender effects on Stretch Attendance Posture (D), in 68 F1 rats from a reciprocal cross between strains RHA and RLA. x-axis indicates direction of reciprocal cross (fRHA/mRLA *vs* fRLA/mRHA) in A-C and sex (M: Male, F:Female) in D. y-axis is behavioral phenotype score (A: Time in Open Section/s, B: number of Entries into Open Section, C: Latency to enter into an open section/s, in Elevated Zero Maze). Thick black horizontal line indicates median, grey box indicates range of central 50% of data points, and outer whiskers the 90% range.

In the elevated zero maze there were no significant differences between fRHA/mRLA or fRLA/mRHA rats in numbers of head dips or defecation, irrespective of gender. Gender was significant in stretch attend postures (SAP); females performed significantly more SAP (P=0.00075) (Figure 3D), but there was no SAP effect for PoE (P= 0.409) and no significant interaction between these factors. There were significant PoE for time spent in the open sections of the maze (P=0.00306), the number of entries (P=0.00367) and the number of line crossings (P=0.01728).

In a two-way active avoidance in the shuttle box no significant differences between the fRHA/mRLA and fRLA/mRHA rats were observed for escape/avoidance latency, inter trial crossings, number of avoidances and context-freezing. There was also no gender effect or interaction between gender and parental origin.

## DISCUSSION

In this study we have surveyed the heritability in HS rats of 199 complex traits, 86% of which show an apparent PoE. Our results are broadly consistent with an extensively-phenotyped population of HS mice, in which we also identified pervasive PoE in the heritability of complex traits ^7^. In HS mice, the enrichment for PoE, as measured by the median ratio of 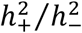 was smaller than reported here in HS rats (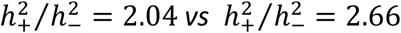 respectively). However, the smaller HS rat sample means that the estimates have larger standard errors (average 0.133 for 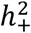 and 0.172 for 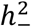 in rats compared to 0.058 and 0.078 in mice). We did not attempt to map PoE QTLs, because of low power due to the relatively small sample size ^7^. These estimates are based on genome-wide genetic similarities, and so are unlikely to be driven entirely by variation at imprinted genes. Whilst confounding with shared environment might explain some of the excess heritability, our results suggest that PoE also originate from loci that are not imprinted. In addition to shared environment, it should be noted that dominance can also generate apparent PoE^11^. It is therefore important to verify the PoE we detected by heritability analysis of HS using an orthogonal experimental design such as reciprocal cross, as we have done here.

There are two differences between the mouse and rat HS experiments requiring comment. First, the HS mice were housed in cages with 4-5 same-sex littermates. The HS rats were housed in cages with just two same-sex littermates. This meant that in the mice, it was possible to remove cage effects before estimating heritability, but in the rats it was not possible to do this as they were too closely confounded. Consequently the estimates of PoE heritability in the rats are likely to be inflated by shared environment. Second, the rats were more closely related than the mice in the sense that many of the parents were cousins. This alters the distribution of allele sharing attributed to parents of opposite sex (*K*^−^, Fig 1B) and makes the distributions of *K*^+^, *K*^−^ more alike in the rats than the mice, thereby potentially reducing our power to detect differences between PoE and non-PoE heritability

The phenotypes obtained from the HS rats included a large battery of behavioral measures. These included avoidance acquisition in the shuttle box and context-conditioned freezing ^26^ both of which assess coping style, and which is also predicted by and correlated with latency to the first entry into an open section and entries to the open section of the elevated zero maze ^25,26,31,51,53–55^.

We observed PoE in 9/10 coping style behaviors in HS rats, which we confirmed in a reciprocal cross of the Roman low-and high-avoidance rats. These strains exhibit opposite responses to stress ^50^: RLA and RHA rats have respectively reactive and pro-active coping styles^29,31,54^, while HS rats have a profile similar to the RLA rats in their coping style and responses to stress ^53,54^. We found F1 rats with RHA fathers and RLA mothers behaved in the zero maze with an RLA-like reactive coping style, whilst those with RLA fathers and RHA mothers more closely resembled RHA rats. These differences remained significant even after removing litter effects, so having a phenotypic response linked to the parental strain cannot be attributed solely to shared environment or maternal effects. Epigenetic modifications in the serotonin and glutamate receptors within in the prefrontal cortex differ between RLA and RHA rats, supporting the general principle that PoE might affect behaviour via epigenetic effects in these strains ^58^. Besides differing in coping style, RHA and RLA rats exhibit different impulsivity, behavioral inhibition and behavioral flexibility. Thus RHA vs RLA differences align with those in reciprocal crosses of mice in which the imprinted genes *Gbr10* or *Nesp55* are knocked out^40,41^.

Thus, as is thought to happen in humans with the same traits ^39,59^, RHA rats present functional deficits in the prefrontal cortex and hippocampus and amygdala, three very closely linked structures that are involved in coping, impulsivity, behavioral flexibility and behavioral inhibition. They also present differences in the serotoninergic system (and other neurotransmitters) with respect to RLA rats. Our results therefore suggest that it would be worthwhile to investigate the impact of PoE on human behaviour.

## ACKNOWLEDGEMENTS

This work was partially supported by the Wellcome Trust core grant 090532/Z/09/Z (to C.M, P.H.P, J.F, R.M.), and by grants PSI2013-41872-P and PSI2017-82257-P (MINECO), 2014SGR-1587(DGR) and “ICREA-Academia2013” (to A.F-T).

---

**Supplemental Table 1**. PoE heritabilities. The table contains the traits, their heritabilities for PoE (“H2.POE“) and additive genetic effects (“H2.NOPOE“) and the standard errors, as well as the number of individuals measured per trait (N). The columns “H2.PATERNAL”, “H2.MATERNAL”, “SE.PATERNAL”,”SE.MATERNAL” contain the paternal and maternal heritabilities and their standard errros. The Type of trait is either N (non-behavioral), CS (coping-style), or B (behavioral excluding coping-style).

**Supplemental Table 2**

Zero Maze Phenotypes from a RLA/RHA reciprocal cross. Each row corresponds to a single rat. Columns are: cage (id of the cage), mstrain (strain of the mother), mID (identifier for mother), fstrain (strain of the father), fID (identifier for father), sex, (sex of animal), ID (individual id), zlatency (time/s to first entry into the open sections of the zero maze), timeopen (time/s spent in open sections of the maze), entries (number of entries into maze open sections), Head_Dips (number of head dips through the edge of the open sections of the maze), SAP (number of stretch attend postures from a closed to an open section of the maze), LineX (number of line crossings in the elevated zero maze).

**Supplemental Table 3**

Shuttle Box Phenotypes from a RLA/RHA reciprocal cross. Each row corresponds to a single rat. Columns are: cage (id of the cage), mstrain (strain of the mother), mID (identifier for mother), fstrain (strain of the father), fID (identifier for father), sex, (sex of animal), ID (individual id), Freezing5 (context freezing, i.e. time spent freezing during the first 5 inter-trial intervals of the two-way avoidance-shuttle box-training session), avoid (number of avoidances during the 40-trial avoidance session), latency (time/s to escape/avoid, averaged for the 40 training trials of the avoidance session), ittx (inter-trial crossings, during the forty 60-min inter-trial intervals of the avoidance session).

